# Retinal outputs depend on behavioural state

**DOI:** 10.1101/638049

**Authors:** Sylvia Schröder, Nicholas A. Steinmetz, Michael Krumin, Marius Pachitariu, Matteo Rizzi, Leon Lagnado, Kenneth D Harris, Matteo Carandini

## Abstract

The operating mode of the visual system depends on behavioural states such as arousal^1,2^. This dependence is seen both in primary visual cortex^3–7^ (V1) and in subcortical brain structures receiving direct retinal input^4,8^. Here we show that this effect arises as early as in the output of the retina. We first measured activity in a region that receives retinal projections^9^, the superficial superior colliculus (sSC), and found that this activity strongly depended on behavioural state. This modulation was not mediated by feedback inputs from V1 as it was immune to V1 inactivation. We then used Neuropixels probes^10^ to record activity in the optic tract, and we found some retinal axons whose activity significantly varied with arousal, even in darkness. To characterize these effects on a larger sample of retinal outputs, we imaged the activity of retinal boutons^11,12^ in sSC during behaviour using a calcium indicator. The activity of these boutons correlated with arousal as strongly as that of sSC neurons, and this correlation persisted also during darkness. These results reveal a novel property of retinal function in mice, which could be observed only during behaviour: retinal outputs are modulated by behavioural state before they reach the rest of the brain.

---

Behavioural states related to the level of arousal or locomotion influence many parts of the brain^1,2^. In the visual system, this influence can increase or decrease neurons’ firing rates^4–8^, affecting both spontaneous activity and visually driven responses^3,4,7^. The effects of behavioural state could be mediated by neuromodulatory input and may involve feedback from higher cortical areas^2^. Once these effects are established at a given stage in the visual pathway, one would expect them to propagate to subsequent stages. For instance, the effects of arousal on neurons in lateral geniculate nucleus^4^ are likely to pass on to the next stage, primary visual cortex (V1). This raises an intriguing hypothesis: that behavioural states may affect the output of the very first stage of visual processing, the retina. Behavioural state might affect the activity of retinal neurons through the inputs that the retina receives from the rest of the brain^13,14^. Behavioural state, moreover, might also affect retinal synapses in their target area, through presynaptic neuromodulation^15^. To test this hypothesis, however, one needs to measure retinal outputs with neuronal resolution during behaviour, a measurement that has traditionally posed substantial challenges.

To measure the effects of arousal in a region targeted by the retina, we imaged the activity of neurons in superficial superior colliculus (sSC) and found strong effects of arousal (Figure 1). We placed mice on a treadmill where they could run while head-fixed (Figure 1a), and assessed arousal by running speed and pupil size^1^. To image sSC without damaging the brain, we developed a custom implant (Figure 1b,c). This implant accesses posterior sSC, where neurons represent the lateral and upper periphery of the visual field (Figure 1d). We expressed the calcium sensitive fluorescence indicator GCaMP6f^16^ non-selectively in sSC neurons via a viral vector, and recorded neuronal activity with two-photon imaging (Figure 1e). Arousal strongly modulated neural responses to drifting gratings (Figure 1f). Correlation with running speed^8^ and with pupil diameter was positive in some neurons and negative in others, and were readily visible in the activity of single neurons (Figure 1f). Across >3,000 sSC neurons, 43% had significant correlations (28% positive, 15% negative) with pupil diameter during visual responses (p < 0.05, shift test; Figure 1g,h). Similar results were seen during spontaneous activity (Figure S1a-c) and when gauging arousal by running speed (Figure S1d-g). We ensured that these correlations were not due to brain movement by simultaneously imaging the activity-independent fluorescence of tdTomato (see Methods).

**Figure 1.**
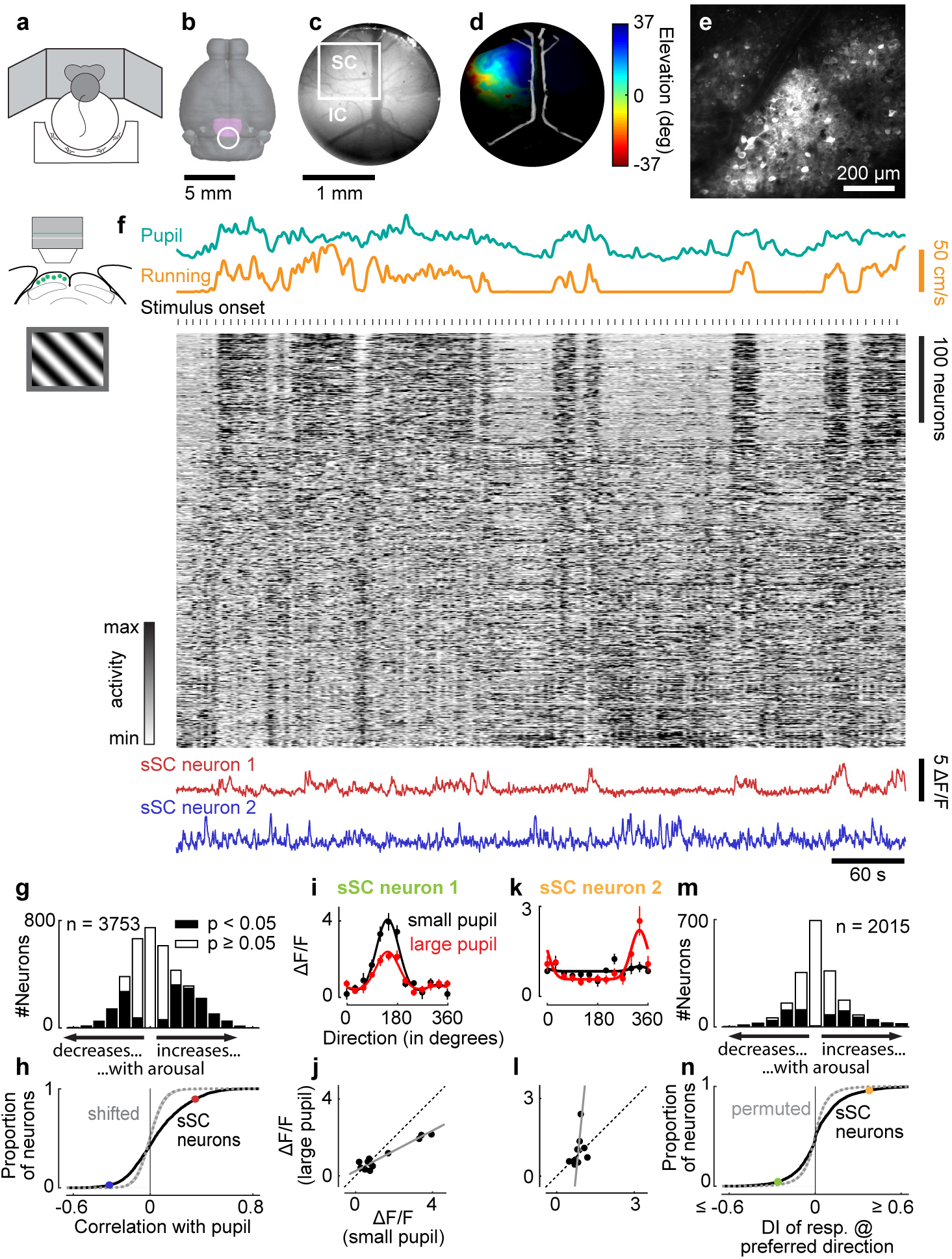
Arousal modulates activity in sSC. **a**, Mice were head-fixed on a treadmill and surrounded by three monitors. **b**, Top view of mouse brain showing SC (purple) and position of implant (circle). **c**, View through implant showing SC and inferior colliculus (IC) and field of view of a two-photon imaging session (white rectangle). **d**, Retinotopic map (via intrinsic optical imaging) of visual elevation in left SC (same brain as in c). Brightness represents signal-to-noise ratio. **e**, Average frame of two-photon imaging data shown in f. **f**, Pupil diameter (green), running speed (yellow), calcium traces of sSC neurons (each z-scored, then sorted by correlation with pupil diameter on first half of data, only second half is presented to show the robustness of the correlations), and traces of two neurons (red and blue) during presentation of gratings. Each grating was presented for 2 s with 3-5 s of grey screen between trials. **g**, Correlation strengths of sSC neurons with pupil diameter during presentation of gratings. Significance of each cell was assessed by random time-shifting. **h**, Cumulative distribution of correlations for data in g (solid line) and randomly time-shifted null distribution (dotted line and shaded area are median and 2.5-97.5 percentile interval). Dots show correlation strengths of two example neurons from f. **i**, Direction tuning (mean±SEM) of sSC neuron measured when pupil was small (black) or large (red). Solid lines show fitted tuning curves. **j**, Scatterplot of responses during small pupil versus during large pupil (same data as in i). Solid line is linear fit derived from fitted tuning curves (i; slope < 1, p < 0.05, permutation test). **k,l**, Same plots as in i,j for another sSC neuron (in l, slope > 1, intercept < 0, p < 0.05, permutation test). m, Distribution of change in response to preferred direction during small versus large pupil, quantified as difference index DI = (L-S)/(L+S), where L and S are responses to preferred direction during large and small pupil. **n**, Cumulative distribution of data in m (solid line) and of time-shifted data (dotted line and shaded area, median and 2.5-97.5 percentile interval). Dots show values of two neurons in i,k.

Consistent with earlier observations^1,3,8^, arousal had additive and multiplicative effects on visual responses, without changing their selectivity for orientation and direction (Figure 1i-l; see Methods for test of linearity across population). To quantify the effects of arousal on each neuron, we measured the responses at the preferred direction (or in mean response, for untuned neurons). Arousal increased these responses in 17% of neurons and decreased them in 15% (p < 0.05, permutation test; Figure 1m,n). In neurons that were selective for orientation or direction, we also measured the tuning depth, i.e. the difference between response to preferred and non-preferred direction. Arousal decreased this measure in 20% of sSC neurons and increased it in 10% (p < 0.05, permutation test; Figure S2).

These effects of arousal in sSC were not mediated by top-down inputs from primary visual cortex (V1) (Figure 2). V1 projects to sSC directly^17^ and influences visual responses of sSC neurons^18^. Given that arousal modulates V1 responses, this modulation might be passed on from V1 to sSC. To test this hypothesis, we recorded activity in superior colliculus (SC) while silencing optogenetically^19,20^ the retinotopically matched part of V1 on half of all trials (Figure S3). These experiments were performed in PV-Cre × Ai32 mice, which express ChR2 in cortical parvalbumin-expressing inhibitory neurons, and SC activity was recorded using electrophysiology, which is more easily combined with optogenetics. When V1 was active, visual responses to preferred direction increased with arousal in 27% of neurons and decreased in 9% (p < 0.05, permutation test; Figure 2a-c), consistent with the results we had seen with two-photon imaging (Figure 1m,n). Inactivating V1 on average decreased the visual responses in SC neurons^18^ independent of the level of arousal (p < 0.05, Wilcoxon signed rank test; Figure 2g,h; difference index (I-C)/(I+C) where I is maximum response to gratings when V1 inactivated and C is maximum responses during control condition: −0.06±0.02 for small pupil and −0.05±0.01 for large pupil). V1 inactivation did not, however, diminish the effects of arousal on SC neurons: arousal increased responses at preferred direction in 25% of SC neurons and decreased them in 13% (p < 0.05, permutation test; Figure 2d-f). In fact, arousal affected responses to preferred stimuli slightly more during V1 inactivation (p = 0.02, Wilcoxon signed rank test; Figure 2i,j), and only 2% of SC neurons were affected more strongly by arousal in control conditions than during V1 inactivation (p < 0.05, permutation test). Effects of arousal on tuning depth of SC neurons did not change significantly with V1 inactivation (Figure S4). Thus, the effects of arousal on SC neurons cannot be explained by inputs from V1.

**Figure 2.**
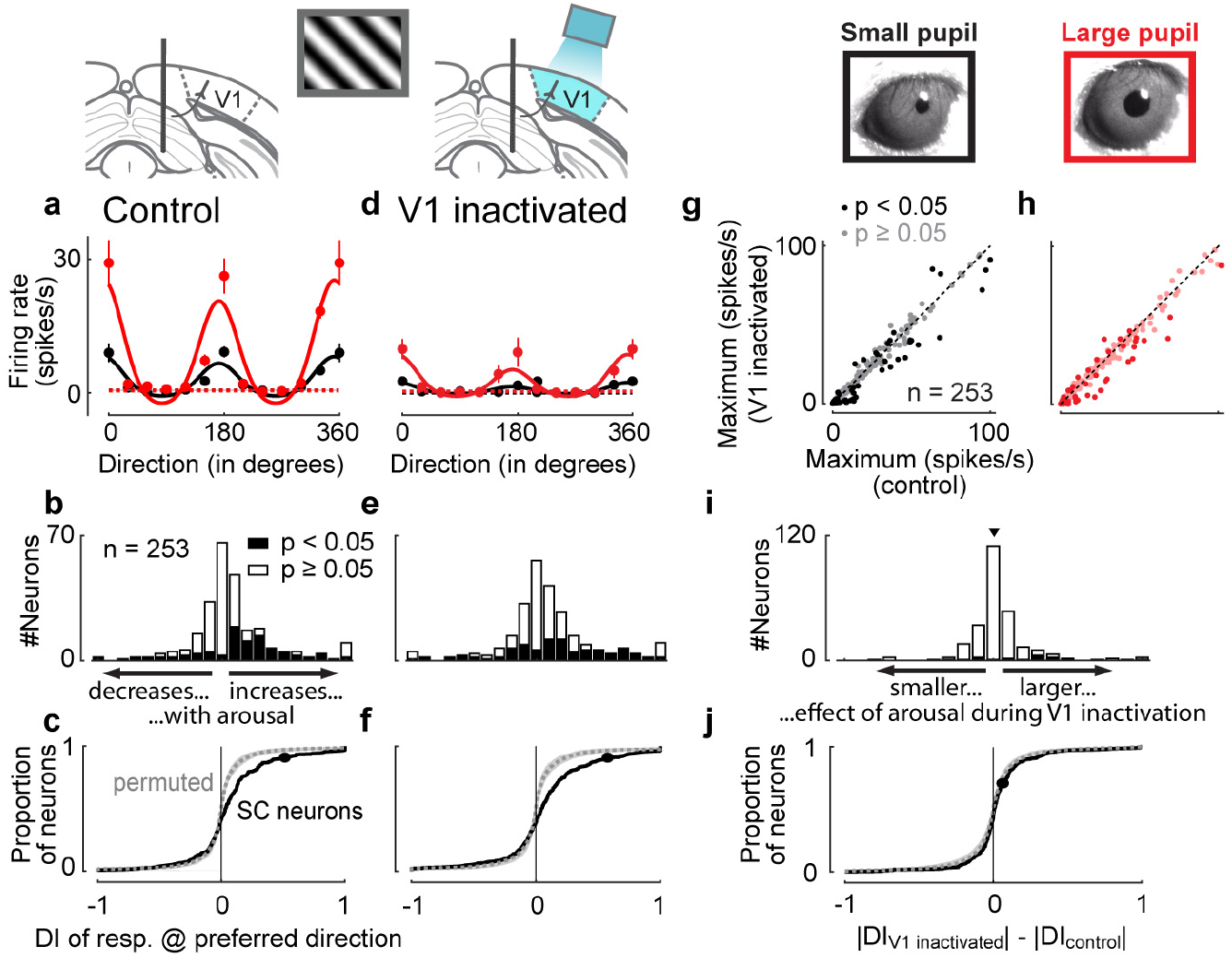
Effect of arousal in SC is not mediated by V1. **a**, Direction tuning (mean±SEM) of SC neuron during small (black) and large (red) pupil. Solid lines show fitted tuning curves. Dotted lines show baseline firing rates. **b,c**, Distribution of change in response to preferred direction during small versus large pupil, quantified as difference index DI (see Figure 1m showing same measure for sSC neurons recorded with two-photon imaging). Dotted line and shaded area in c show median and 2.5-97.5 percentile interval of permuted data. Black dot in c shows value for example neuron in a. **d**, Direction tuning of same neuron as in a during inactivation of V1. **e,f**, Same plots as b,c but during V1 inactivation. **g,h**, Fitted maximum tuning responses of all SC neurons during control condition vs. V1 inactivation. Pupil was either small (g) or large (h). Darker dots: cells whose, maximum response changed significantly during V1 inactivation (p < 0.05, permutation test). i,j, Comparison of arousal effects between control condition and V1 inactivation. Distribution shows difference between |DI| during V1 inactivation and |DI| in control condition. Median difference (triangle in i) is 0.006 (p = 0.02, Wilcoxon signed rank test). Dotted line and shaded area in j show median and 2.5-97.5 percentile interval of permuted data. Black dot in j shows value for example neuron in a,b.

To assess whether arousal modulation might be found in retinal inputs, we recorded the activity of single retinal axons in the optic tract (Figure 3a-d). We recorded with Neuropixels probes^10^ and adopted a spike sorting algorithm that can track the spikes of single units if the brain moves relative to the electrode during behaviour, followed by manual curation. The selection of units was based on stringent criteria. The first criterion was anatomical: units were selected if histological reconstructions^21^ placed them putatively in the optic tract (Figure S5a,c). The second criterion was visual: units were selected if they gave reliable responses to rapidly flickering stimuli (Figure S5b,d) or had a clear spatial receptive field (Figure 3a,b) and short latency. The third criterion was electrophysiological: units were selected based on spike amplitude and lack of correlation between this amplitude and firing rate. The first criterion yielded 1,280 putative optic tract units, the second narrowed this sample to 49 units, and the third to 25 units. To avoid biases, all of these analyses were performed before considering any measurement of locomotion or arousal. We then considered these measurements, and found, reassuringly, that running caused no changes in spike shape in any of the 25 retinal axons (Figure 3c,d). We thus turned to the effects of arousal on neural activity.

**Figure 3.**
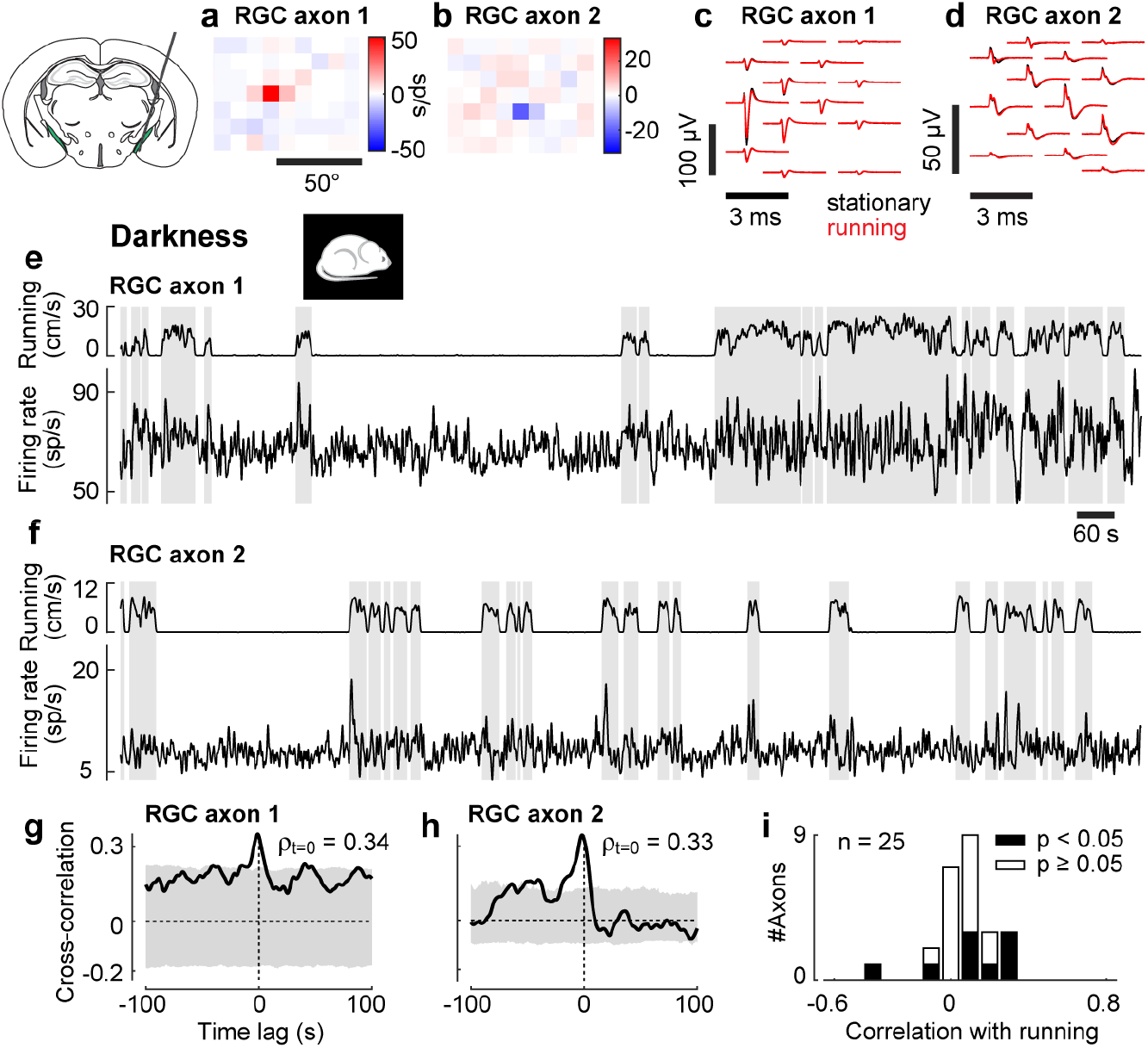
Effect of arousal is present in firing rates of retinal ganglion cells. **a,b**, Receptive field of axon 1 (a) and axon 2 (b) relative to the centre screen. **c,d**, Spike waveform of axon 1 (c) and axon 2 (d) on multiple neighbouring channels of probe when animal was running (red) or stationary (black). Spikes were recorded during darkness. **e,f**, Trace of running speed (top) and firing rate (bottom) of axon 1 (e) and axon 2 (f) recorded in darkness. Grey shades indicate periods of running (≥1 cm/s). **g,h**, Cross-correlogram between firing rate and running speed for axon 1 (g) and axon 2 (h). A positive lag denotes that firing rate is lagging running speed. Grey shade shows 2.5-97.5 percentile interval of time-shifted data. **i**, Correlation strengths (measured at a lag of zero seconds) of all recorded retinal axons with running speed during darkness.

Arousal significantly modulated the activity of several retinal axons, even in darkness (Figure 3e-i). In darkness the pupil is fully dilated, so it cannot reflect changes in arousal. We thus gauged arousal from running speed, which is closely related to pupil size (Figure S1g). Consider the firing rates of two retinal axons recorded during darkness (Figure 3e,f). The cross-correlation between firing rate and running speed shows positive correlations around a lag of zero seconds; these correlations were larger than expected by chance as ascertained by time-shifting the running trace (Figure 3g,h). Positive correlations were seen in 7 of the 25 recorded retinal axons, negative correlations in 2 of 25 axons (p < 0.05, shift test, Figure 3i), more than expected by chance (p = 7.6e-10, Fisher’s combined probability test). Arousal therefore affected the firing rate of retinal axons, and this effect could not be explained by changing light levels (e.g. due to pupil dilation), because these measurements were performed in darkness.

To confirm these effects of arousal on a larger sample of retinal outputs, we expressed a calcium indicator in retinal ganglion cells^11^, and imaged the activity of their synaptic boutons in sSC during behaviour (Figure 4a-e). To target the expression of calcium indicator to the axonal boutons^12^, we cloned a variant of GCaMP6f fused with a localisation signal targeting synaptic terminals^22^ (SyGCaMP6f) and packaged it into a AAV2 vector. We injected this AAV2-SyGCaMP6f into the vitreous humour of one eye (Figure 4a-c). We then performed two-photon imaging in the contralateral sSC (Figure 4d). We spaced the imaging planes very tightly, with <2 μm separation, so that we could track activity of the boutons even if the brain moved perpendicular to the imaging planes (Figure 4e). Because the indicator was localized to synaptic boutons, fluorescence was not affected by fibres of passage.

**Figure 4.**
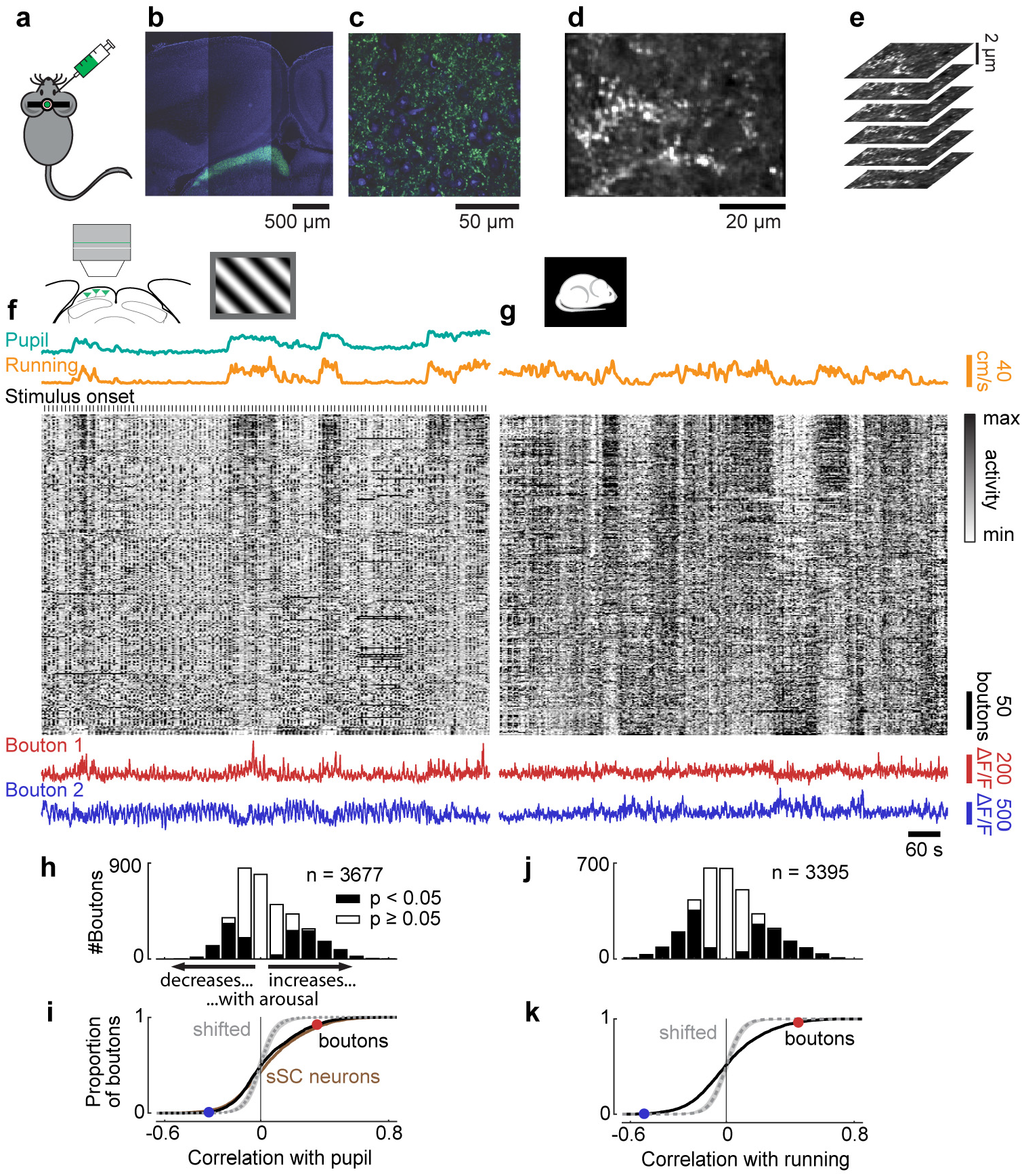
Effect of arousal on activity of retinal boutons in sSC. **a**, SyGCaMP6f is injected into eye contralateral to imaged SC. **b**, Confocal image showing expression of SyGCaMP6f in the sSC. c, Zoom-in to view in b. **d**, Average frame of two-photon imaging data shown in f,g. **e**, Proximal planes were imaged near simultaneously to track boutons during brain movement. **f**, Pupil diameter trace (green), running speed trace (yellow), raster of calcium traces of retinal boutons (grey; each z-scored and vertically sorted by correlation with running speed including of stimulation and darkness), and traces of two boutons (red and blue) during presentation of gratings. **g**, Similar to f but during darkness. Pupil was fully dilated (trace not presented). Raster shows responses of same boutons in same order as in f. **h,i**, Correlation strengths of retinal boutons with pupil diameter during presentation of gratings. For comparison, correlation strengths of sSC neurons are plotted with brown line (i, same data as in Figure 1h). **j,k**, Correlation strengths of retinal boutons with running speed during darkness. Circles in i,k show the correlation strengths of the two boutons in f,g.

Retinal boutons in sSC were strongly modulated by arousal, to a similar extent as sSC neurons themselves (Figure 4f-k). Activity in retinal boutons was strongly correlated with pupil diameter and running speed (Figure 4f). During the presentation of gratings, 41% of all imaged retinal boutons had significant correlations (23% positive, 18% negative) with pupil diameter (p < 0.05, shift test; Figure 4h). The distribution of correlation strengths was similar to that observed in sSC neurons (Figure 4i). When presenting grey screens, correlations with pupil size were similar to those during presentation of gratings, and the distribution of correlation strengths was again similar to that found in sSC neurons (Figure S6a-c). Even in complete darkness, activity of retinal boutons was markedly affected by arousal (Figure 4g) and almost half (45%) of all retinal boutons were significantly correlated (23% positively, 22% negatively) with running speed (p < 0.05, shift test; Figure 4j,k). Correlations with running speed measured during darkness resembled those seen during the presentation of gratings (Figure S6d). Like in sSC neurons, the effects of arousal on direction tuning curves of retinal boutons were linear (Figure S7a-d, see Methods for test of linearity), although unlike SC neurons, boutons showed a consistent bias towards negative modulation by arousal (response at preferred direction: increased in 5% of boutons, decreased in 26%; tuning depth: increased in 3% of tuned boutons, decreased in 28%; p < 0.05, permutation test; Figure S7e-h).

Behaviour could affect retinal outputs through several mechanisms. One potential mechanism are the projections that the retina receives from the rest of the brain^13,14^. In rodents, these projections include histaminergic fibres from hypothalamus and serotonergic fibres from the dorsal raphe^23^. Both neuromodulators reflect behaviour and arousal^24–26^. Additional mechanisms could operate at retinal boutons in superior colliculus, e.g. through presynaptic modulation of calcium influx^15^ by neuromodulators such as serotonin^27,28^.

Modulation of retinal activity across behavioural states might optimize encoding of visual stimuli, for example by controlling the operating range of retinal ganglion cells^23^. Theories of efficient and predictive coding suggest that the retina is attuned to the statistics of natural scenes and to the behavioural needs of the animal^29–31^. Both of these factors change when the animal moves or becomes aroused, and the retina may accommodate this change by adapting its encoding strategy.

## Acknowledgments

We thank Charu Reddy for help with mouse husbandry; we thank Miles Wells, Laura Funnell, Rakesh Raghupathy for help with histology; we thank Jack Waters and Pedro Garcia da Silva for help with the design of the implant used for two-photon imaging; we thank Paul Johnson and Ian Macartney for help with the development of surgical tools; we thank Robin Ali for generous support.

This work was supported by the People Programme (Marie Curie Actions) of the European Union’s Seventh Framework Programme (FP7/2007-2013) under REA Grant Agreement No 62387 (to SS), by the Biotechnology and Biological Sciences Research Council (grant BB/P003273/1 to MC and SS), by the European Union’s Marie Skłodowska-Curie program (fellowship 656528 to NAS), by the Human Frontier Sciences Program (fellowship LT001071/2015-L to NAS), by the Wellcome Trust (grants 205093 and 102264 to MC and KDH; grant 102905/Z/13/Z to LL), and by HHMI Janelia (MP). MC holds the GlaxoSmithKline / Fight for Sight Chair in Visual Neuroscience.

## Author Contributions

S.S., M.C., and K.D.H. conceived and designed the study. S.S. and N.A.S. collected the data. S.S. analysed the data. S.S., N.A.S., M.K., and M.P. wrote code to analyse the data. L.L. designed the SyGCaMP6f transgene. M.R. generated the final AAV2-SyGCaMP6f vector. S.S., M.C., and K.D.H. wrote the manuscript.

## Methods

All procedures were conducted in accordance with the UK Animals Scientific Procedures Act (1986) under personal and project licenses released by the Home Office following appropriate ethics review.

### Animals

We used 32 mice: 11 inbred C57Bl/6J (www.jax.org/strain/000664; 1 male, 10 female) and 21 transgenic mice (13 female, 8 male) were used in this study. For two-photon imaging, we used 15 mice obtained by crossing Gad2-IRES-Cre (www.jax.org/strain/010802) and Ai9 (www.jax.org/strain/007909). The heterozygous offspring expressed TdTomato in glutamate decarboxylase 2-positive (GAD2+) cells to identify inhibitory neurons. For optogenetic inactivation of V1, we used 5 mice obtained by crossing PV-Cre (www.jax.org/strain/008069) and Ai32 (RCL-ChR2(H134R)/EYFP, www.jax.org/strain/012569). The heterozygous offspring expressed ChR2 in parvalbumin-positive cells. For widefield imaging, we used one heterozygous mouse resulting from crossing Emx1-IRES-Cre (www.jax.org/strain/005628) and Ai38 (www.jax.org/strain/014538). Because this mouse did not express GCaMP in SC, we measured intrinsic signals. Animals were 6-41 weeks old at the time of surgery with a mean weight of 27.0 g (19.3-51.6 g) and were used for experiments up to the age of 54 weeks. Mice were kept on a 12-h light: 12-h dark cycle. Most animals were single housed after the first surgery.

### Surgical procedures

Animals were anesthetized with isoflurane (Merial) at 3.5% for induction, and 1-2% during surgery. Carprofen (5 mg/kg; Rimadyl, Pfizer) was administered subcutaneously for systemic analgesia, and dexamethasone (0.5 mg/kg; Colvasone, Norbrook) was administered as an anti-inflammatory agent to prevent brain swelling. The scalp was shaved and disinfected, and local analgesia (Lidocaine, 6 mg/kg, Hameln pharmaceuticals ltd) was injected subcutaneously under the scalp prior to the incision. Eyes were covered with eye-protective gel (Chloramphenicol, Martindale Pharmaceuticals Ltd). After the animal was placed into a stereotaxic apparatus (5% Lidocaine ointment, TEVA UK, was applied to ear bars), the skin covering and surrounding the area of interest was removed, and the skull was cleaned of connective tissue. A custom made headplate was positioned above the area of interest and attached to the bone with Superbond C&B (Sun Medical). Throughout all surgical procedures, the animal was kept on a heating pad to stabilize body temperature at 37°C. Subcutaneous injections of 0.01 ml/g/h of Sodium Lactate (Hartmann’s solution) were given. After the surgery, the animal was placed into a heated cage for recovery from anaesthesia. Mice were given three days to recover while being treated with carprofen.

In animals used for two-photon imaging, a circular 4 mm craniotomy (centred at approximately −4.2 mm AP and 0.5 mm ML from Bregma) was made using a biopsy punch (Kai medical) and a fine-tipped diamond drill (Type 250-012 F, Heraeus). To reduce bleeding from the bone and from the dura we used bone wax and gel foam, and we cooled the area by applying cold cortex buffer. As the posterior SC is covered by a large sinus running through the dura, we permanently pushed the sinus anteriorly to gain optical access to the SC. We first made a cut into the dura directly posterior to the transverse sinus spanning the whole width of the craniotomy. Then we inserted a custom-made implant into the cut and pushed it anteriorly and a few 100 microns down to apply some pressure on the brain and thus stabilize the imaging. The implant was made of a short tube (2.29 mm inner diameter, 1 mm length) made of stainless steel (MicroGroup, Medway, Massachusetts). A 3 mm glass coverslip (#1 thickness, World Precision Instruments) was glued onto the tube to seal the end that was inserted into the craniotomy. A stainless steel washer was glued onto the other end of the tube. The washer had an inner diameter that fit exactly around the tube and an outer diameter of 5 mm (Boker’s, Minneapolis, Minnesota). All three pieces were glued to each other using a UV curing adhesive (NOA61, Thorlabs). The glass coverslip was slightly larger than the outer diameter of the tube so that it could be slipped underneath the dura. The implant was placed into the craniotomy so that the washer was sitting on top of the skull and provided stability for the implant. The implant was fixed to the skull with Vetbond (3M) and Superbond C&B (Sun Medical). To prevent any dirt from staining the glass coverslip, we filled the tube of the implant with Kwik-Cast (World Precision Instruments), which could be easily removed before imaging.

For two-photon calcium imaging of activity in sSC neurons, we injected the virus AAV2/1.Syn.GCaMP6f. WPRE.SV40 at a final concentration of 2.30-4.39e12 viral particles/ml after making the cut into dura. 115-230 nl of the virus was injected 300-500 μm below the brain surface at the posterior edge of the right SC, which abuts the inferior colliculus. The virus was injected at a rate of 2.3 nl every 6 s (Nanoject II, Drummond). The injection pipette was kept in place for about 10 min after the end of the injection.

To image retinal boutons in sSC, we engineered a virus to express a calcium indicator localised to synaptic boutons, AAV2-SyGCaMP6f, and we injected the virus into the left eye. We cloned a variant of GCaMP6f fused with a localisation signal targeting synaptic terminals^22^ (SyGCaMP6f) and packaged it into an AAV2/2 vector (Addgene, 51085). This virus restricted GCaMP expression to boutons^12^. To deliver the virus, the animal was anesthetized with isoflurane (Merial) at 3.5% for induction and 1-2% during the procedure. Carprofen (5 mg/kg; Rimadyl, Pfizer) was administered subcutaneously for systemic analgesia. Pupils were dilated by applying a drop of Mydriacil (Alcon, Surrey, UK) into the eye. We then applied Viscotears (Alcon) onto the eye and placed a 3-5 mm glass coverslip over the eye to allow for good visual access to the retina. Using a Hamilton syringe (5 μl; needles: 34 gauge, RN NDL, 10 mm length, point 4), we injected approximately 3 μl of AAV2/2.hSyn1.SyGCaMP6f.SV40 with a concentration of 2.44 × 10^12^ viral particles/ml into the vitreous humour.

For electrophysiological recordings, craniotomies were performed above the areas of interest (anterior SC and V1, or optic tract). For large craniotomies (3 mm diameter) we increased stability by inserting a honeycomb disk^32^ in the craniotomy. The disk was made of polycarbonate with a 3 mm diameter and 175 μm thickness, and had 19 holes of 500 μm diameter each in a hexagonal pattern (laser cut by Laser Micromachining Ltd, Denbighshire, UK). The disk was fixed to the bone with superglue (Powerflex, Loctite).

### Two-photon imaging

Two-photon imaging was performed using a standard resonant microscope (B-Scope, ThorLabs Inc.) equipped with a 16×, 0.8 NA water immersion objective (N16XLWD-PF, Nikon) and controlled by ScanImage 4.2 (Ref. ^33^). Excitation light at 970-980 nm was delivered by a femtosecond laser (Chameleon Ultra II, Coherent). Multi-plane imaging was performed using a piezo focusing device (P-725.4CA PIFOC, Physik Instrumente, 400 μm range). Laser power was depth-adjusted and synchronized with piezo position using an electro-optical modulator (M350-80LA, Conoptics Inc.). The imaging objective and the piezo device were light shielded using a custom-made metal cone, a tube, and black cloth to prevent contamination of the fluorescent signal caused by the monitors’ light. Emission light was collected using two separate channels, one for green fluorescence (525/50 nm emission filter) capturing the calcium transients and one for red fluorescence (607/70 nm emission filter) capturing the expression of TdTomato in inhibitory neurons of Gad-Cre × TdTomato mice.

For imaging neurons in sSC, we used 3-4 imaging planes separated by 9-30 μm at depths of 15-100 μm from the surface of SC. The field of view spanned 340-640 μm in both directions at a resolution of 512 × 512 pixels. The frame rate per plane was 6.0-7.5 Hz. One dataset captured a single plane at a resolution of 1024 × 1024 pixels, and with a frame rate of 15 Hz.

For imaging retinal boutons in sSC we used 5 or 10 imaging planes (with 3 or 6 fly-back planes) with an interplane distance of <2 μm at depths of 8-46 μm from the surface of SC. The field of view spanned 75-135 μm at a resolution of 256 × 256 pixels, or 42-10 μm at a resolution of 128 × 128 pixels. The frame rate per plane was 7.5 Hz.

### Intrinsic widefield imaging and retinotopic map

To obtain the retinotopic map in Figure 1d, we performed intrinsic imaging using methods described previously^34^. Presentation of stimuli (periodic moving bars) and data analysis were as described in the paper.

### Eletrophysiology

Recordings were made using Neuropixels electrode arrays^10^. Probes were mounted to a custom 3D-printed piece and affixed to a steel rod held by a micromanipulator (uMP-4, Sensapex Inc.). To allow later track localization, prior to insertion probes were coated with a solution of DiI (ThermoFisher Vybrant V22888 or V22885) or DiO (ThermoFisher Vybrant V22886) by holding 2 ⊠l in a droplet on the end of a micropipette and touching the droplet to the probe shank, letting it dry, and repeating until the droplet was gone, after which the probe appeared pink. Probes had a soldered connection to short external reference to ground; the ground connection at the headstage was subsequently connected to an Ag/AgCl wire positioned on the skull. The craniotomies and the wire were covered with saline-based agar. The agar was covered with silicone oil to prevent drying. In some experiments a saline bath was used rather than agar. Probes were advanced through the agar and the dura, then lowered to their final position at ~10 ⊠m/sec. Electrodes were allowed to settle for ~10 min before starting recording. Recordings were made in external reference mode with LFP gain = 250 and AP gain = 500. Data were filtered in hardware with a 300 Hz 1-pole high pass filter and digitized at 30 kHz. Recordings were repeated at different locations on each of multiple subsequent days.

During recordings with optogenetic inactivation of V1, one electrode was placed into SC and a second electrode into V1. In 5 of 6 datasets, both electrodes were recorded simultaneously. The SC electrode entered the brain vertically at −3.7 mm AP and ±0.6 mm ML from Bregma. The V1 electrode entered the brain at −3.6 (or −3.5) mm AP and ±2.25 (or +3.0) mm ML from Bregma. This electrode was tilted backward from vertical by 20° and tilted away from the midline by 45°. 4 of 6 recordings were performed in the right hemisphere.

For recordings from the optic tract, the electrode entered the brain at −1.7 (or −2.0) mm AP and ±2.62 (or −1.6) mm ML from Bregma. The electrode was tilted backward from vertical by 10° and tilted away from the midline by 90° so it was aligned with coronal plane of the brain. 3 of 6 recordings were performed in the right hemisphere.

### Optogenetic inactivation

For optogenetics experiments, 473 nm light was generated by a diode laser (LuxX, Photon Lines Ltd.), coupled into an optic fiber, and then focused to a spot ~1 mm in diameter on the surface of the brain near the probe. Light intensity was modulated in a 40Hz raised cosine wave pattern, with peak light power at the surface of the brain of approximately 0.9-3 mW. The exact light power for each experiment depended on how strongly neural activity simultaneously measured in V1 was affected. The optogenetic inactivation was performed on for 700 ms including 100 ms before and after the visual stimulus. In one dataset, the inactivation laser was switched on and off simultaneously with the visual stimulus, which was presented for 2 s.

### Experimental setup and visual stimuli

The mouse was head fixed with a headplate holder that did not obstruct the visual field. For two-photon imaging, the mouse was free to run on an air-suspended Styrofoam ball (20 cm in diameter), whose rotation was measured by two optical computer mice^35^. For electrophysiology, the mouse was free to run on a Styrofoam wheel (15 cm wide, 18 cm diameter), whose rotation was measured by a rotary encoder (1,024 pulses per rotation, Kübler, Germany). The mouse was acclimated to head-fixation for at least three days before the first recording session, step-wise increasing fixation time from 10 min on the first day to 1 h on the last day.

The mouse was surrounded by three computer screens (Iiyama ProLite E1980SD placed ~20 cm from the mouse’s eyes; or Adafruit, LP097QX1 placed ~11 cm from the mouse’s eyes; 60 Hz refresh rate for both models) at right angles covering approximately 270 × 70 degrees of visual angle. In some experiments, Fresnel lenses (BHPA220-2-6 or BHPA220-2-5, Wuxi Bohai Optics) were mounted in front of the monitors to compensate for reduction in luminance and contrast at steeper viewing angles relative to the monitors. In some of these experiments, lenses were coated with scattering window film (frostbite, The Window Film Company) to prevent specular reflections. To track the eye contralateral to the recording site (except in 5 of 7 recordings in SC during V1 inactivation and 3 of 6 recordings in optic tract), we illuminated the eye with an infrared LED (850 nm, Mightex SLS-0208-A or Mightex SLS-0208-B). Videos of the eye were captured at 30 Hz with a camera (DMK 23U618 or DMK 21BU04.H, The Imaging Source) equipped with a zoom lens (Thorlabs MVL7000) and a filter (long-pass, Thorlabs FEL0750; or band-pass, combining long-pass 092/52×0.75, The Imaging Source, and short-pass FES0900, Thorlabs).

Grating stimuli were presented full-field at 100% contrast. Gratings were sinusoidal with spatial frequency of 0.08 cycles per degree and drifting with temporal frequency of 2 cycles per second. Gratings were presented for 2 s separated by a grey screen for 3-6 s. Only in some electrophysiological recordings in SC, the duration of the gratings was 0.5 s with an inter-trial interval of 0.5-1.0 s. The stimuli were repeated 15 or 20 times. During two-photon imaging, the red channel of the monitors was switched off to reduce light contamination in the red fluorescence channel.

To map receptive fields, we presented checkerboard images with white, black and mean grey squares with an edge length of 10 visual degrees (in one dataset 4 visual degrees). The stimulus updated at a rate of 6 Hz and was presented for 10 min. In each noise image, each square was randomly assigned its luminance value with a 98% probability of being grey and 1% probabilities of being black and white.

Spontaneous activity was recorded either when screens were grey or during darkness. Grey screens were presented at mean luminance of the monitors. The duration was 10 min for sSC neurons and 5 or 10 min for retinal boutons. For recordings during darkness, all monitors were switched off and other light sources in the recording rig were eliminated or covered. The recording setup was enclosed either by a frame tightly covered with black curtains or by a closed box, in order to prevent light in the recording room entering the eyes of the animal. The duration of the recordings were 5, 10, or 20 min for retinal boutons and around 40 min for optic tract axons (data used for correlation analyses was 11.7-43 min long; see below for selection of valid time periods).

### Perfusion and Histology

Mice were perfused with 4% PFA, the brain was extracted and fixed for 24 hours at 4 C in PFA, then transferred to 30% sucrose in PBS at 4 C. The brain was mounted on a microtome in dry ice and sectioned at 60 ⊠m slice thickness. Sections were washed in PBS, mounted on glass adhesion slides, and stained with DAPI (Vector Laboratories, H-1500). Images were taken at 4× magnification for each section using a Zeiss AxioScan, in three colours: blue for DAPI, green for DiO, and red for DiI.

### Data analysis

#### Preprocessing of two-photon imaging data

All raw two-photon imaging movies were analysed using Suite2p to align frames and detect regions of interest^36^. We used the red channel representing TdTomato expressed in all inhibitory neurons to align frames, which yielded better results than alignment using calcium dependent fluorescence. For 22 of 28 datasets, alignment was non-rigid. In the remaining ones rigid alignment was sufficient.

For imaging data of retinal boutons, we also aligned frames in depth, i.e. when the brain moved perpendicular to the imaging planes, fluorescence data from neighbouring imaging planes was used to correct this movement. This was possible because imaging planes were very close to each other (<2 μm) so that fluorescence from any neural structure was detected in multiple planes. First, the target images of all planes (except fly-back planes) were aligned to each other in x and y. Then each frame of the red channel data was aligned to each target image and the similarity to each target image was determined. For each imaging cycle through all planes, the optimal shift of this stack of images was determined so that the similarity of all planes to the target image of their assigned target planes was maximal. The shifted planes were then assigned to the corresponding target plane. For each imaging stack, a moving average across neighbouring planes (width 2-4 planes depending on similarity between planes) was applied so that neighbouring planes became more similar to each other. A fluorescence trace containing values from neighbouring planes across time thus resulted in a smooth trace. The same alignment and averaging steps were also applied to frames of the green channel, using offsets estimated from the red channel. Boutons were analysed only if they appeared in every cycle, meaning that they never moved outside the imaged volume. The algorithm of alignment throughout depth is implemented in Suite2p. ROIs were detected using the aligned frames of the green channel, and were then manually curated using the Suite2p GUI.

All aligned movies were inspected manually to check for failures in automatic alignment. Failures were corrected using different parameter settings where possible. Otherwise, the misaligned movie frames were discarded (1.1±0.6% of frames per dataset were discarded).

Using the aligned movies and detected ROIs resulting from Suite2p analysis, we extracted the fluorescence from the green and the red channel within each ROI. To correct the calcium traces for contamination by surrounding neuropil, we also extracted the fluorescence of the surrounding neuropil for each ROI using the green channel. The neuropil mask resembled a band surrounding the ROI with its inner edge having a distance of 3 microns away from the edge of ROI (for neurons; 1 micron for boutons) and its outer edge having a distance of 30 microns from the edge of the ROI. Pixels belonging to other ROIs were excluded. To correct for contamination, the resulting neuropil trace, N, was subtracted from the calcium trace, F, using a correction factor α: F_c_(t) = F(t) - α∙N(t). The correction factor was determined for each ROI as follows. First, F and N were low-pass filtered using the 8^th^ percentile in a moving window of 180 s, resulting in F_0_ an N_0_. The resulting traces F_f_(t)=F(t)-F_0_(t) and N_f_(t)=N(t)-N_0_(t) were then used to estimate α as described previously^37^. In short, N_f_ was linearly fitted to F_f_ using only time points when values of F_f_ were relatively low and thus unlikely to reflect neural spiking. F_c_ was then low-pass filtered as above (8^th^ percentile in a moving window of 180 s) to determine F_c,0_. These traces corrected for neuropil contamination were then used to determine ΔF/F = (F_c_(t) - F_c,0_(t)) / max(1, mean_t_(F _c,0_(t)).

To correct for potential brain movements, we used the red fluorescence traces of each ROI to regress out changes in fluorescence that were not due to neural activity. First, we low-pass filtered the red trace of each ROI (8^th^ percentile in a moving window of 180 s) and subtracted it from the unfiltered trace to remove slow drifts and bleaching effects. Second, we applied a median filter to the resulting red trace (moving median in window of 10 s). Third, this trace was regressed out of ⊠F/F.

To avoid counting the same cell or bouton multiple times due to its detection in multiple imaging planes, we detected ROI pairs with highly correlated calcium traces (⊠ > 0.4 for neurons and ⊠ > 0.5 for boutons, correlation between traces filtered using a moving median in a window of 5 samples) and nearby ROI centres in neighbouring imaging planes. Only the ROI of each pair with the highest signal-to-noise ratio was used for further analyses. ROIs that had very long-lasting calcium transients (>25 s) were removed.

#### Spike sorting

Extracellular voltage traces were preprocessed using common-average referencing: subtracting each channel’s median to remove baseline offsets, then subtracting the median across all channels at each time point to remove artefacts. Electrophysiological data collected in SC was spike sorted using Kilosort with standard parameters^38^. Data collected in the optic tract was spike sorted using a modification of Kilosort, termed Kilosort 2 (code available at http://www.github.com/MouseLand/Kilosort2). Kilosort 2 is able to track spikes of a neuron when its location relative to the probe changes, i.e. during drift. In addition, Kilosort2 performs automated splits and merges similar to what a human curator would do on the basis of spike waveform similarity, the bimodality of the distribution of waveform features, and the spike auto- and cross-correlograms. After sorting, all automatically detected spike clusters were curated manually using Phy (github.com/kwikteam/phy).

#### Criteria for selection of retinal axons in optic tract

The first step in identifying spiking units that correspond to axons in the optic tract was the histological analysis of the brains. Using the traces of DiI or DiO left behind by the probe, we determined whether the probe passed through the optic tract, and if so, which part and thus which recording sites were located in the optic tract. Using DAPI staining, the optic tract is readily identifiable so that recordings that did not pass the optic tract could easily be discarded. If the probe did pass through the optic tract, we next used SHARP-Track^21^ to align each brain slice to a plane through the Allen Mouse Common Coordinate Framework (Allen Institute for Brain Science, 2015) and record the 3D coordinates of manually selected points along the fluorescence track. A line was fitted through the coordinates, resulting in a vector of brain regions the electrode passed through. Next we localized the position of the probe along this vector of brain areas including the matching scaling factor for this vector. In most cases, it was not possible to locate the tip of the probe in the brain slices with precision high enough to align probe position with the vector of brain areas; instead we used physiological features varying along the probe, such as spike rate and spike amplitude, to match those to the identified brain areas along the reconstructed track^32^. For example, the lower edge of cortical cell layers and the different cell and dendritic layers of hippocampus were often readily identifiable from the recordings. After this alignment, we selected those recording sites that were estimated to pass through the optic tract (36 of 50 recordings).

The second step in identifying optic tract units was the screening for visual responses indicative of retinal axons. The probe was inserted into the brain so that brain areas recorded on sites close to the optic tract were not part of the early visual system, e.g. internal capsule and globus pallidus above, and medial amygdala below optic tracts. These areas were not expected to have strong visual responses with short latencies. We presented three different visual stimuli to test those visual response properties. The stimuli were composed of (1) luminance reversing (flickering) screens with reversal rates of up to 15 Hz to measure visual responsiveness to fast changes in luminance, (2) visual noise consisting of white and black squares on grey background to measure receptive fields, and (3) full-field drifting sinusoidal gratings to measure direction tuning. Only units that showed a clear visual response and temporal modulation to the fastest flickering screens (15 or 7.5 Hz) were considered to be retinal axons. For very few units, no responses to flickering monitors were collected or the firing rate in response to these stimuli were extremely low. In these cases, units were included only if they had well-defined receptive fields with shapes and sizes expected of retinal ganglion cells, short response latencies to grating stimuli (<50 ms), and unusual spike waveforms (for example like those in Figure 3d). Of the 1,280 units that were located in the estimated position of the optic tract, 49 units passed our criteria for retinal axons.

The final step of selecting optic tract units was to discard those units that did not pass our criteria for stable and reliable recordings. Because retinal axons are very fine structures electrophysiological recordings from them are prone to artefactual changes in firing rates resulting from electrode drift. The use of densely spaced recording sites on Neuropixels probes (nearest site at most 20 μm apart^10^) together with a spike sorting algorithm that automatically tracks recorded units across electrode drift (see above) helped us to account for the consequences of electrode drift. Still, we employed additional criteria to control that no artefacts contaminated the data used to study the correlation between firing rates and running speed. To avoid biases, we did not consider the recorded running speed when applying these criteria. First, low-amplitude clusters were excluded: if the lowest amplitude spikes in the cluster touched the detection threshold at any point, the entire cluster was not analysed (18 of 49 neurons were excluded by this criterion), or the affected time periods of the unit were discarded. Second, any units with a significant correlation between firing rate and spike amplitude were excluded as potentially contaminated by electrode drift (5 of 49 neurons excluded; significance of correlation was determined using the shift test as described below in Correlation analyses).

If after automatic spike sorting the distribution of spike amplitudes was bimodal, we identified and discarded the small amplitude contaminating spikes using an automatic method. To do so, spikes were binned so that each bin contained 100 spikes. For each time point, the most frequent spike amplitude, termed mass amplitude, was determined. The vector of mass amplitudes was smoothed (moving average on 3 bins) and all spikes with amplitudes above the current mass amplitude were used to estimate the SD of the distribution of spike amplitudes (for each spike, the mass amplitude at spike time was subtracted; the resulting distribution of positive amplitudes was fit with a half-Gaussian). Any spikes with amplitudes smaller than the respective mass amplitude minus 5-20 SDs (varying between units) were discarded from further analysis. This was done in 10 of 26 units (for 7 units during darkness). Finally, one of these 10 units had to be excluded because the mouse hardly ran during the remaining periods in darkness (<5%). We thus included 25 units in our correlation analysis.

We then took into account the recorded running speeds of the animal and plotted for each unit its mean spike waveform during a running episode and a temporally nearby stationary episode. We checked that spike waveforms did not change between the two episodes, which would point to electrode movement (see Figure 3c,d for mean spike waveforms during running and stationarity pooled across all spikes of 2 example neurons recorded during darkness). No further units had to be excluded based on this analysis.

#### Tracking of pupil

Movie processing was performed offline using custom code written in MATLAB (Mathworks) on a frame-by-frame basis. Briefly, each frame was mildly spatially low-pass filtered to reduce noise and then the pupil contour was detected by a level-crossing edge detector. Then, concave segments (caused by the reflections of the infrared LED or by whiskers) of the estimated pupil contour were discarded, and the position and the area of the pupil were calculated from the ellipse fit to the concave segments of the pupil contour. Eye-blinks were detected by a two-dimensional classifier, based on overall frame intensity (frames with blinks tend to be brighter) and correlation to the average frame (frames with blinks tend to have lower correlation to the average frame). The output of the algorithm was visually inspected, and adjustments to the parameters (e.g., spatial filter strength, level-crossing threshold, boundaries of the blink detection classifier) were made if necessary. The trace of pupil size was then smoothed by using the median in a moving window of 5 samples. Frames with detected blinks were excluded from the analysis.

#### Correlation analyses

For two-photon imaging experiments, traces of running speed and pupil diameter were interpolated to match the sampling rate of the calcium traces. All signals were then convolved with a Gaussian window with a sigma of 1 s. Correlation strength was determined using Pearson’s correlation. Because both neural activity and pupil/running are temporally autocorrelated, significance was tested by time shifting. A null distribution was obtained by circularly shifting the running or pupil trace by a random amount of time, and then calculating Pearson’s correlation with the calcium traces, 500 times. The correlation of the non-shifted data was significantly different from zero if it fell below the 2.5 percentile or above the 97.5 percentile of the null distribution. Note that for display purposes, the raw traces of the two example neurons in Figure 1f and two example boutons in Figure 4f,g were only smoothed with a moving average over 5 samples (0.67 s).

For electrophysiological recordings from retinal axons in the optic tract during darkness, spikes were binned into bins of 133 ms (to match the commonly used sampling rate of 7.5 Hz for calcium imaging), and running traces were interpolated to samples at the same time points. Running traces and firing rates were then convolved with Gaussian window with sigma of 1 s, after slow drifts were subtracted (drift was estimated by 8^th^ percentile in moving window of 180 s). Then the cross-correlogram of the z-scored signals were calculated. If the signals were composed of separate parts of the data (i.e. when certain parts were excluded by our quality criteria measures), cross-correlograms were calculated separately on each part and then averaged. Significance was tested as above by a shift test. If the signals were composed of separate parts, these were concatenated before the shifts.

#### Fisher’s combined probability test

To verify whether the number of cells correlating with a variable exceeded that expected from independent chance, we used Fisher’s method of combined probabilities. If ρ_c_ represents the p-value for a test of cell *c*, Fisher’s method combines these into a chi-square statistic 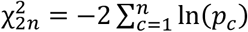, which follows a 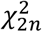 distribution under the assumption that all p-values are independent.

#### Estimation of responses to gratings

From calcium traces recorded in an sSC neuron or a retinal bouton, we quantified response magnitudes to drifting gratings by iteratively fitting a temporal response kernel that is 15 s long and a scaling factor for each presentation of a grating. First, we used a General Linear Model (not to be confused with Generalized Linear Model) to fit one temporal kernel to all trials using the currently estimated scaling factors (initially set to 1). Second, we used the currently estimated temporal kernel and fitted a scaling factor for each trial, again using a General Linear Model. Both steps were repeated until the values of the estimated scaling factors stopped changing across iterations. We used the scaling factors as response strength in each trial.

The reason for using this approach rather than stimulus-triggered averages of the calcium trace was the dynamics of the calcium decay, which in some neurons was so slow that it extended into the response to the subsequent stimulus. Our kernel fits correct for these calcium residues in stimulus responses and represent the neural responses very well (adjusted R^2^: 0.4±0.007). We only included neurons that showed significant responses to the gratings. To test significance we randomly shifted the response of each neuron against the stimulus presentation times. A neuron was determined to be *responsive* if the mean square error (MSE) of the fits to the measured responses was smaller than the 95% confidence interval of MSEs resulting from fits to the randomly shifted responses, which was the case for 2015 of 3753 detected neurons.

To estimate response amplitudes from electrophysiological recordings, we calculated firing rates from spikes between stimulus onset and offset.

#### Fitting tuning curves

Two different tuning curves were fitted for responses during small and large pupil trials. To categorize each trial, the trace of pupil size was interpolated to match the sampling times of the neural data (spikes were binned into 5 ms bins) and then thresholded at the median pupil size measured during the experiment. If pupil size was below that threshold for longer than half of the trial, the trial was categorized as small pupil trial, and otherwise as large pupil trial.

Before fitting, we inverted the responses of all suppressed-by-contrast neurons, i.e. neurons whose activity mostly dropped in response to a full-field grating stimulus. Note that for these neurons their preferred direction elicited the largest drop in activity.

Responses during the same pupil size were fit to two wrapped Gaussians with peaks separated by 180 degrees. Each curve had five parameters: preferred direction, response at preferred direction (on top of offset), tuning width (sigma of Gaussian), direction selectivity ((P- N)/(P+N), where P is response to preferred direction, N is response to null direction), and offset. Tuning width (sigma) was limited to a minimum of the sampling distance between tested grating directions. All curves were fitted using the least-squares method.

Tuning curves for small and large pupil were constrained to have equal preferred direction, tuning width, and direction selectivity. First, one tuning curve was fit to all responses independent of pupil size to find good initial parameters for the following fit. Then two tuning curves were for the two pupil sizes under the condition that the three above parameters are the same for both curves.

During experiments involving V1 inactivation, we fitted two tuning curves during control conditions and two tuning curves during V1 inactivation. The only parameter that was fixed across all four curves was preferred direction, which was determined by first fitting all responses to a single tuning curve. The fitting procedure for data during small and large pupil was the same as described above.

To decide whether the neuron was tuned to direction or not, we compared the wrapped Gaussian fit to constant fit across stimuli. The two models were compared using cross-validated explained variance, i.e. the data of one repetition for each stimulus was left out for fitting and predicted responses were then compared to the left-out data (this was repeated for as many times each stimulus was presented). Neurons were considered tuned if the cross-validated fit with the Gaussian resulted in larger explained variance than the constant fit (1308 of 2015).

Responses to preferred direction in untuned neurons were defined as the constant number fitted to responses to all stimuli. Tuning depth (difference between the maximum and minimum fitted response) was only defined for tuning neurons.

To test whether arousal had significant effects on the tuning curve, i.e. whether the tuning curve fit to the low arousal data was significantly different to the tuning curve fitted to the high arousal data, we used a permutation test. We shuffled the arousal categorization across trials and refitted tuning curves on the shuffled data, 200 times. For each shuffling, we computed a test statistic (e.g. difference in mean firing rate to preferred stimulus between aroused and non-aroused states). The measure on the original data was deemed significant if it lay outside the 2.5-97.5 percentile interval of the shuffled data.

Similarly, the effects of V1 inactivation were tested by shuffling laser on and off categorizations and comparing measures on the original data to the 2.5-97.5 percentile interval of the shuffled data.

#### Test for linearity of arousal effect on tuning curves

As described above, we fitted two tuning curves to grating responses during intervals with small and large pupil, with the constraint that both curves have the same preferred direction, tuning width and direction selectivity. The resulting tuning curves can only differ in their additive offset and their gains. These two tuning curves thus define a linear relationship between the small pupil and large pupil responses and determine intercept and slope of the line fit. To test whether the effects of arousal are linear or not, we tested whether the neurons’ responses could be fit better when relaxing our constraints. We calculated cross-validated explained variance as described above for three further models: (1) no constraints, (2) same preferred direction, and (3) same preferred direction and tuning width. We then compared the resulting explained variances from all four models using ANOVA and found no significant differences between the four distributions (p = 0.91, F = 0.18).

To test the significance of fitted intercepts and slopes in Figure 1j,l, we used the same permutation test as described above. We shuffled the arousal categorization across trials and refitted tuning curves on the shuffled data, 200 times. For each shuffling, we computed the fitted intercept and slope. Intercept and slope of the original data were deemed significant if they lay outside the 2.5-97.5 percentile interval of the shuffled data.

#### Mapping of receptive fields

To map the linear receptive fields of the axons recorded in the optic tract, we counted spikes during the presentation of each visual noise image (presented at a rate of 6 images per second) and subtracted the mean spike count across all images. Using a General Linear Model, we then fitted a kernel for each square location of the noise stimulus in response to the appearance of a non-grey square (we did not differentiate between white and black). The kernel spanned the presentation of 6 images (corresponding to a length of 1 s). Figure 3a,b shows the kernels of all squares at the time when the kernel with the maximum absolute value was largest.

---

**Figure S1.**
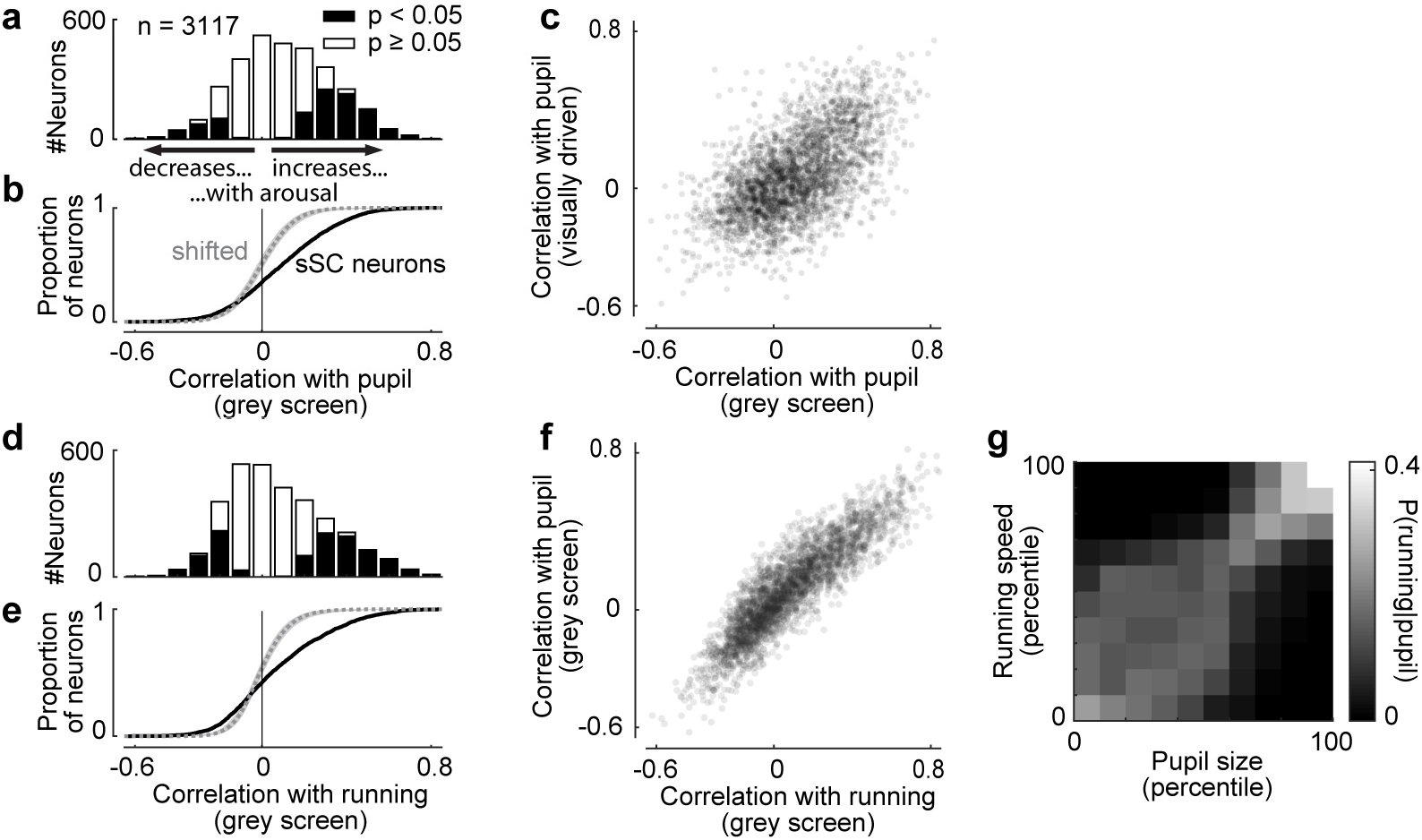
Behavioural effects in sSC neurons generalize across different measures of arousal and across different visual input. **a**, Correlation strengths of sSC neurons with pupil diameter during presentation of grey screens (spontaneous activity). 27% of all neurons had a significantly positive correlation with pupil size, 8% a significantly negative correlation (p < 0.05, shift test). **b**, Cumulative distribution of correlations for data in a (solid line) and time-shifted data (dotted line and shaded area are median and 2.5-97.5 percentile interval). **c**, Correlations with pupil size during presentation of gratings (visually driven) are plotted against correlations with pupil size during presentation of grey screens. Pearson correlation ρ = 0.58 (p < 0.001, Student’s t-test, t = 39.5, df = 3115). **d,e**, Same plots as in a,b but for correlations with running speed during presentation of grey screens. 25% of sSC neurons were positively correlated, 13% negatively (p < 0.05, shift test). **f**, Correlations with pupil size plotted against correlations with running speed. Neural activity was measured during presentation of grey screens. Pearson’s ρ = 0.64 (p < 0.001, Student’s t-test, t = 109.9, df = 3115). Data in a-f were collected from same neurons. **g**, Distribution of running speed (quantified in percentiles of all measured running speeds) given the simultaneously measured pupil diameter (quantified in percentiles). Each column sums to 1. The plot shows that running speed and pupil size strongly depend on each other.

**Figure S2.**
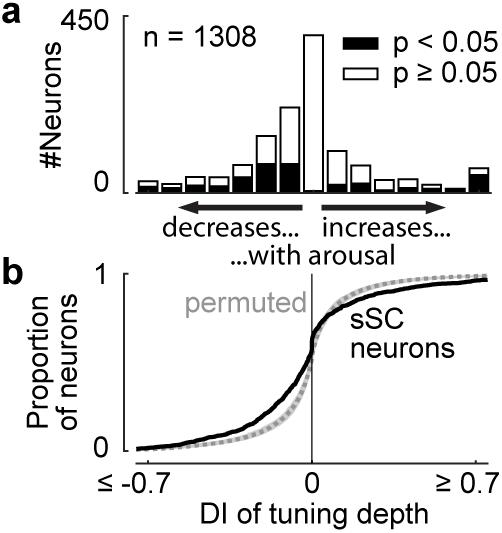
Effects of arousal on tuning depth in sSC neurons. **a**, Distribution of change in tuning depth during small versus large pupil, quantified as difference index DI (see Figure 1m). Only tuned neurons were included here. **b**, Cumulative distribution of data in a (solid line) and permuted data (dotted line and shaded area show median and 2.5-97.5 percentile interval).

**Figure S3.**
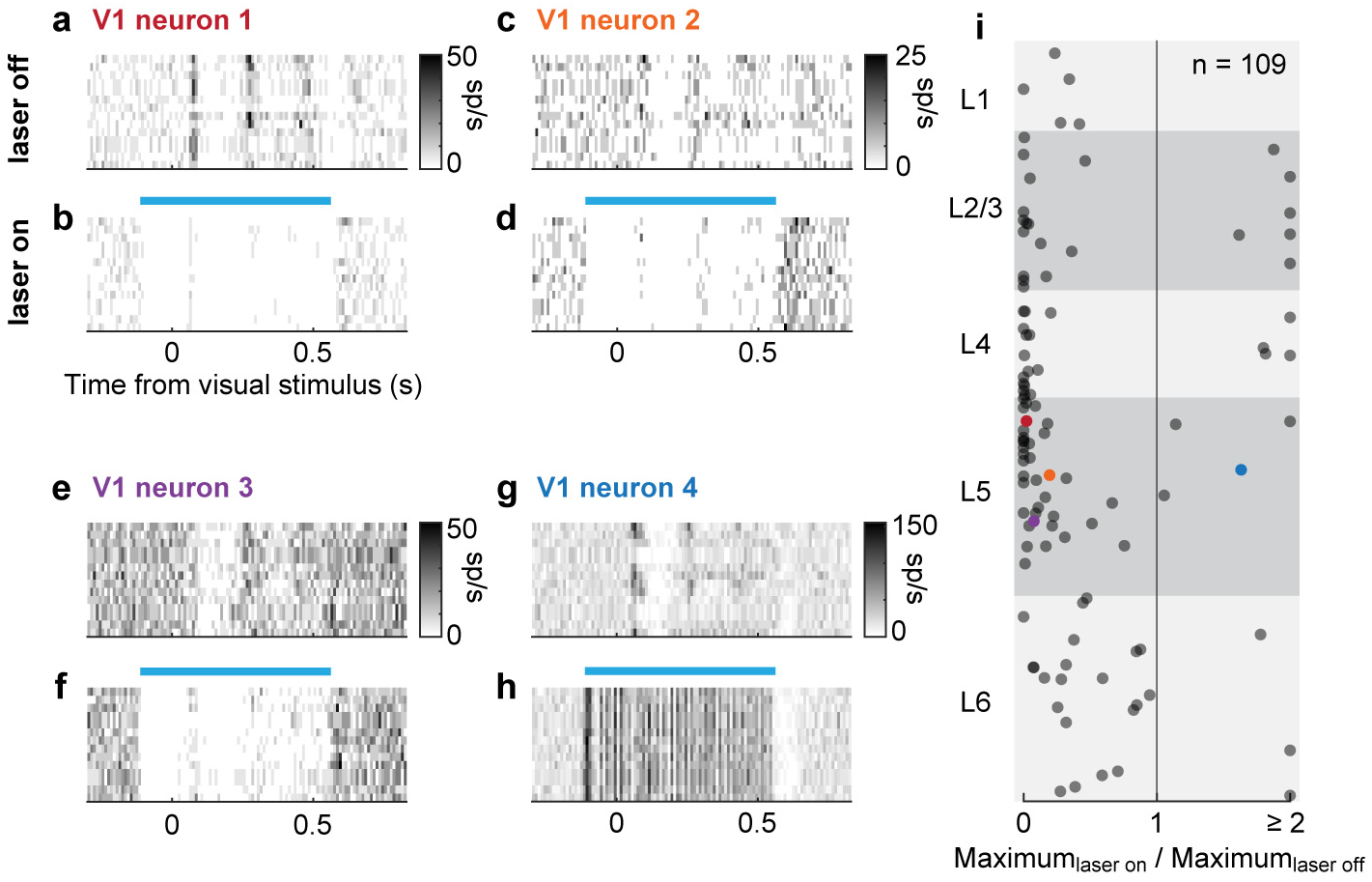
Optogenetic inactivation of V1. **a**, Mean firing rate (colour coded) of a layer 5 (L5) V1 neuron during the presentation of gratings drifting in 12 different directions (each row shows response to a different stimulus). Each grating is presented for 0.5 s. **b**, Same as in A but here the laser was switched on from 0.1 s before the onset of the visual stimulus until 0.1 s after stimulus offset (blue bar). As can be seen from the plot, the neuron was strongly inactivated when the laser was on. **c-h**, Same plots as in a,b for three other L5 V1 neurons. Note that neuron 4 (g,h) was strongly activated when the laser was on and thus is a putative parvalbumin-positive, inhibitory neuron. i, Ratio of visual response to stimulus evoking largest response among all presented gratings (i.e. not fitted) during V1 inactivation and during control condition, plotted against the depth of the neurons within V1. Depth is scaled between 0 (surface of V1) and 1 (bottom of layer 6). Neurons with ratios larger than 1 are putative inhibitory neurons that are excited by laser stimulation. Layer 5 of V1 contains neurons projecting to SC. Coloured dots: example neurons in a-h. Data were collected from two experiments.

**Figure S4.**
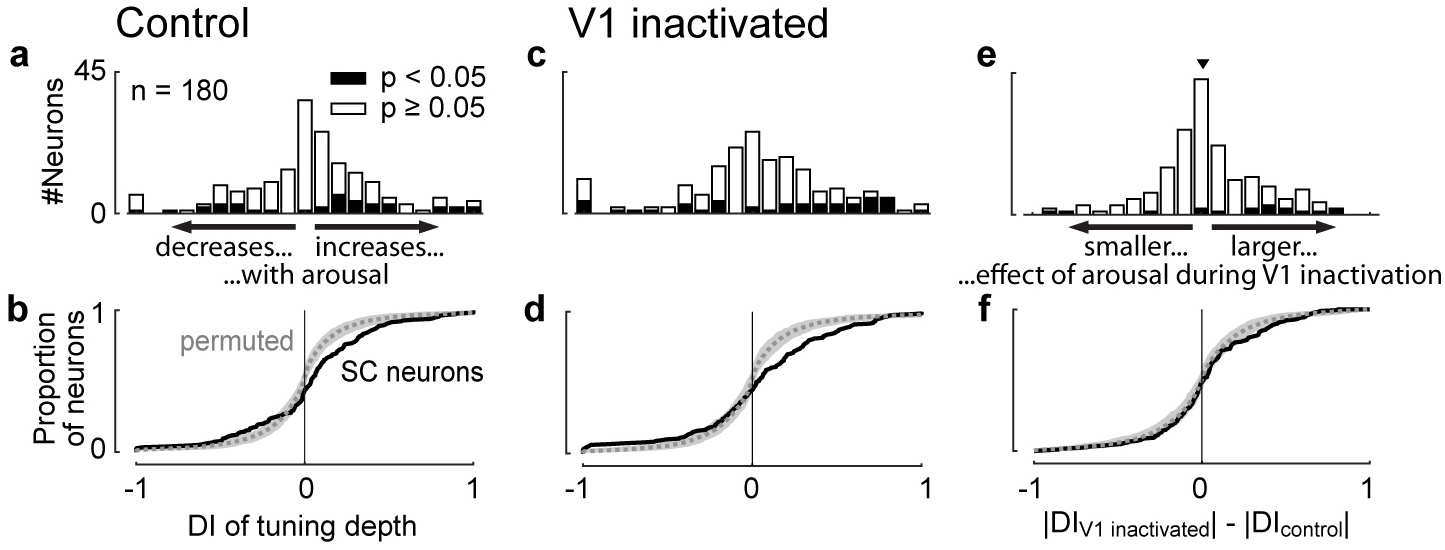
Effect of arousal on tuning depth of SC neurons during control conditions and V1 inactivation. **a,b**, Distribution of change in tuning depth during small versus large pupil, quantified as difference index DI (see Figure 1m). Tuning depth refers to the difference between fitted maximum and fitted minimum response to gratings. 14% of SC neurons increased tuning depth with arousal, 7% decreased (p < 0.05, permutation test). Dotted line and shaded area in b show median and 2.5-97.5 percentile interval of permuted data. **c,d**, Same plots as a,b but during V1 inactivation. 16% of SC neurons increased tuning depth with arousal, 9% decreased (p < 0.05, permutation test). **e,f**, Comparison of arousal effects between control condition and V1 inactivation. Distribution shows difference between |DI| during V1 inactivation and |DI| in control condition. There is no significant difference between absolute DIs in control conditions and during V1 inactivation. Median difference (triangle in e) is 0.01 (p = 0.77, Wilcoxon signed rank test) showing that tuning depth of SC neurons did not change during V1 inactivation. Dotted line and shaded area in f show median and 2.5-97.5 percentile interval of permuted data.

**Figure S5.**
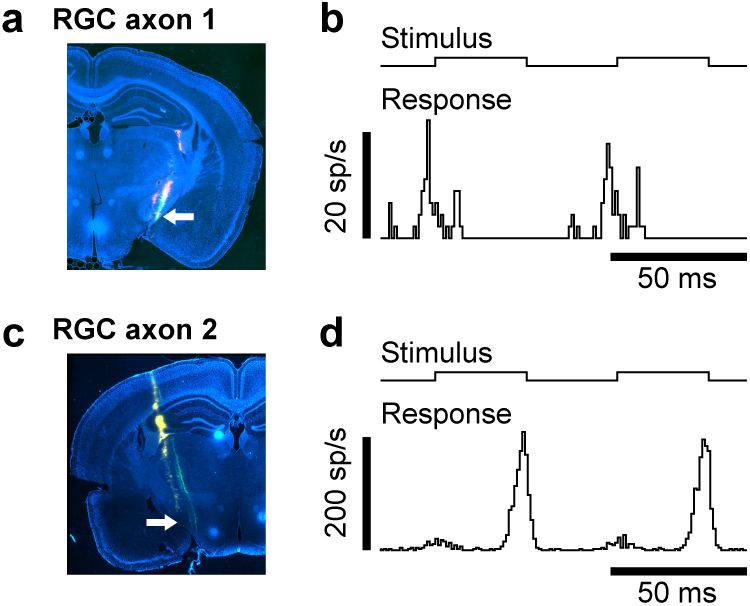
Histology and visual responses of example retinal axons in Figure 3. **a**, Coronal brain slice showing tracks of multiple recordings (green and red). Recording of axon 1 indicated by arrow. **b**, Average response (bottom trace) of axon 1 to presentation of full-field reversing (black/white) stimulus (top trace). Luminance changed every 33.3 ms. c,d, Same plots as a,b for axon 2.

**Figure S6.**
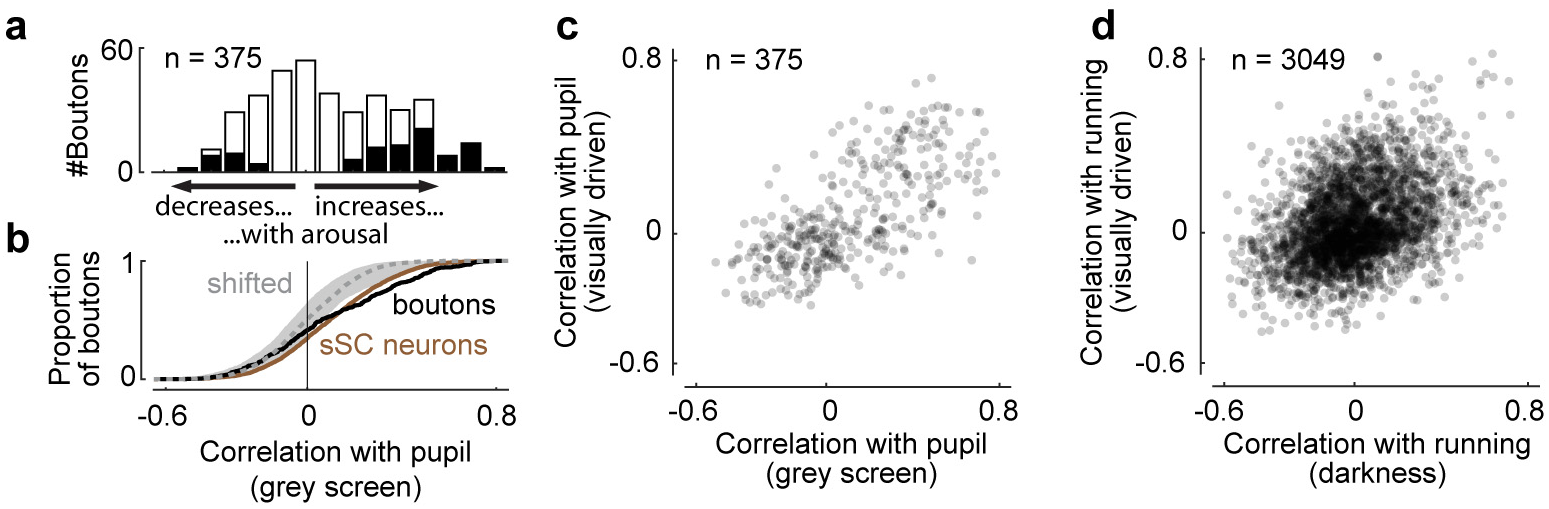
Effect of arousal on spontaneous activity of retinal boutons. **a,b**, Correlation strengths of retinal boutons with pupil diameter during presentation of grey screens (spontaneous activity). 20% of the boutons were positively correlated with pupil size, 6% negatively (p < 0.05, shift test). Dotted line and shaded area in b show median and 2.5-97.5 percentile interval of time-shifted data. For comparison, correlation strengths of sSC neurons are plotted with brown line (same data as in Figure S1b). **c**, Correlations with pupil size during presentation of gratings (visually driven) are plotted against correlations with pupil size during presentation of grey screens. Pearson’s ⊠ = 0.69 (p < 0.001, Student’s t-test, t = 18.3, df = 373). **d**, Correlations with running speed during presentation of gratings (visually driven) are plotted against correlations with running speed during darkness. Pearson’s ⊠ = 0.36 (p < 0.001, Student’s t-test, t = 21.4, df = 3,047).

**Figure S7.**
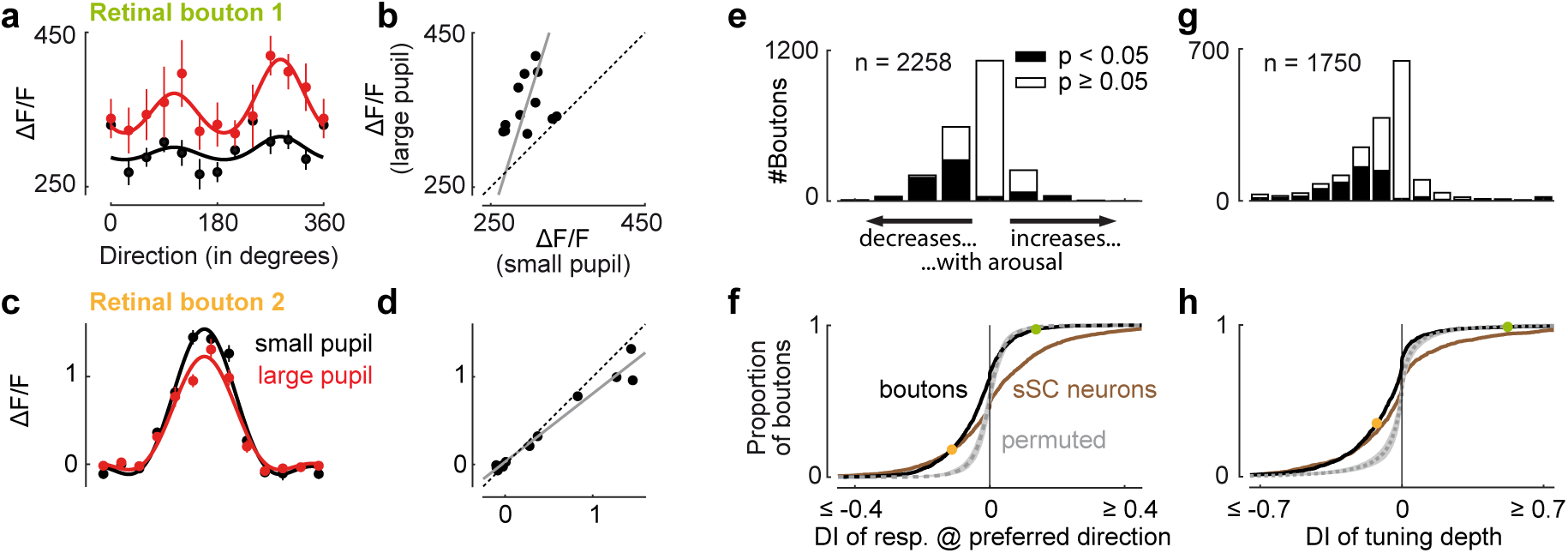
Effect of arousal on tuning of retinal boutons. **a**, Direction tuning (mean±SEM) of a retinal bouton measured when pupil was small (black) or large (red). Solid lines show fitted tuning curves. **b**, Scatterplot of responses during small pupil versus during large pupil (same data as in a). Solid line is linear fit derived from fitted tuning curves (a, slope > 1, intercept < 0, p < 0.05, permutation test). **c,d**, Same plots as a,b for another retinal bouton (in d, slope < 1, p < 0.05, permutation test). **e,f**, Distribution of change in response to preferred direction during small versus large pupil, quantified as difference index DI (see Figure 1m). Dotted line and shaded area in f show median and 2.5-97.5 percentile interval of permuted data. For comparison, DI values of sSC neurons are plotted with brown line (same data as in Figure 1n). **g,h**, Same plots as e,f for DI of tuning depth. Data of sSC neurons are same as in Figure S2b. Dots in f,h show values of two boutons in a,c.

